# Auxetic patch material exhibits systolic thickening and restores pump function in a finite element model of acute myocardial infarction repair

**DOI:** 10.1101/2023.11.02.565400

**Authors:** Joseph Borrello, Zirui Zheng, Ana Cristina Estrada, Kevin D. Costa

**Affiliations:** Mount Sinai BioDesign, Department of Neurosurgery, Icahn School of Medicine at Mount Sinai; Cardiovascular Research Institute, Department of Cardiology, Icahn School of Medicine at Mount Sinai; Department of Biomedical Engineering, Worcester Polytechnic Institute; Department of Biomedical Engineering, Yale University

**Keywords:** Auxetic, Patch plasty, Ventricular support device, Myocardial infarction, Simulation

## Abstract

Passive mechanical reinforcement of the infarcted heart has been shown to counteract infarct expansion and left ventricular (LV) functional degradation. However, traditional patch plasty of the ischemic region also restricts diastolic filling, reducing cardiac output. These negative side-effects can be minimized with strategic modification of the standard patch, suggesting further functional improvements could be possible through a broader exploration of patch materials. This study examines the potential advantages of a patch graft with auxetic properties, having a negative Poisson’s ratio (ν < 0). For preliminary evaluation, an established finite element model of LV biomechanics pre- and post-acute infarction, originally developed for modeling patch plasty with non-auxetic, or meiotic, materials, was modified to simulate epicardial implantation of auxetic patch materials. A homogeneous auxetic patch graft (ν = -0.2) exhibited radial thickening during systole, driven by tension on the patch from the neighboring contracting myocardium; but the patch thickness expanded away from the LV cavity and therefore did not improve ventricular mechanics compared to a standard meiotic patch graft (ν = 0.4). Alternatively, a heterogeneous auxetic patch design with an outer reinforcement layer caused inwardly-directed systolic thickening that restored the LV pressure-volume relationship toward baseline function, without adversely affecting fiber stress in the border zone or remote myocardium compared to the meiotic patch. This computational modeling study demonstrates the potential to harness auxetic mechanical properties for improving LV pump function in the setting of acute myocardial infarction, motivating further experimental validation of auxetic metamaterial patch devices for surgical repair of injured myocardium.

## Introduction

Every year, over half a million Americans suffer a myocardial infarction (MI)^1,2^, characterized by the death or permanent damage of a region of heart muscle caused by the blockage of blood flow to one or more of the coronary arteries that supply the myocardium^3–5^. While acute survival rates have dramatically improved since the 1960s^6–9^, this has led to a growing population living with chronic MI^1,10,11^. This often leads to left ventricular (LV) remodeling that occurs as fibroblasts infiltrate the necrotic region and replace highly organized and contractile cardiomyocytes with a disorganized and non-contractile fibrotic scar^12–14^. As a consequence of LV remodeling, the infarct is stretched and expanded from increased myocardial wall stress as part of a compensatory mechanism to counteract the loss of a contractile region of the heart^15–17^. This biomechanical adaptation, however, can cause a degenerative cycle governed conceptually by Laplace’s Law^18,19^, especially in cases of large, transmural infarcts^20^. The infarct stretching, expansion, and thinning that can result from compensatory hypertrophy and increased wall stress will further reduce cardiac output, leading to ventricular dilation, decompensation, and ultimately, heart failure (HF). MI-induced LV remodeling is now the leading cause of heart failure, which kills approximately 100,000 people in the US each year^2,10,15,21,22^. There is no cure for heart failure; however, as a therapeutic strategy, mechanical unloading of the LV to lower stress on the myocardium has been shown to reduce infarct size and lead to some beneficial effects known as reverse remodeling^23–25^.

Common approaches to facilitate unloading of the LV include direct mechanical interventions to fully or partially support the myocardium. These interventions include active forms of unloading, such as ventricular assist devices (VADs) 26,27, as well as passive strategies, such as cardiac, or ventricular, support devices (CSDs/VSDs) 25. While VADs reduce mechanical burden on the heart by taking on some or most of the pumping function, CSDs and VSDs mechanically support the myocardium by covering the infarct, or in some cases the entire heart, with a rigid or semi-rigid material, typically a woven textile or piece of decellularized tissue. These support patches mechanically couple to the healthy, neighboring myocardium, shielding the infarct zone from systolic wall stress to prevent continued infarct stretching and expansion^25,28–30^. In several preclinical and clinical trials, both whole-heart CSDs and infarct-only VSDs have prevented LV dilation^31–34^. However, the reduction in LV compliance beneficial to the redistribution of systolic wall stress and the arrest of dilation, was also associated with a decrease in diastolic filling and associated loss of stroke volume (SV) and cardiac output ^34–39^.

The simplicity and proven short-term efficacy of CSDs and VSDs makes them an appealing platform to improve upon for the treatment of MI-induced HF, which has been explored in several earlier studies from Holmes and colleagues^34,40,41^. Across several *in silico* and *in vivo* studies, they demonstrated that modifying the structure and mechanics of VSDs can improve cardiac performance in the acute post-MI setting, specifically by introducing slits to retain longitudinal stiffness while increasing circumferential compliance to allow greater LV filling. These studies highlight the potential to improve post-MI cardiac biomechanics through unique design modifications to existing forms of passive mechanical reinforcement.

An ideal VSD would dynamically provide mechanical support throughout the cardiac cycle, becoming compliant during diastole and then thickening and stiffening during systole, similar to the behavior of healthy myocardium. Auxetic metamaterials, whose complex internal structures give them the counterintuitive ability to get thicker when stretched rather than thinner (negative Poisson’s ratio)^42,43^, could offer such capabilities. Furthermore, recent developments in advanced manufacturing have made it increasingly feasible to fabricate auxetic metamaterials with the dimensions and constituent materials relevant to VSD patch plasty procedures^44,45^.

In order to better understand how an auxetic metamaterial VSD (auxVSD) might impact cardiac biomechanics post-MI, we adapted a biomechanical model previously published by Estrada and colleagues^46^ to produce a computational biomechanical screening tool that would allow for the evaluation of VSD patches with a variety of auxetic mechanical properties.

## Materials and Methods

The model originally constructed by Estrada and colleagues was based on results obtained from large animal studies evaluating the impact of anisotropic cardiac patches (that were selectively softened either longitudinally or circumferentially) on heart mechanics and infarct healing^34,40,47^. The geometry for these finite element (FE) models was obtained by segmenting cine MRI images captured during *in vivo* studies of the acutely ischemic canine LV. In total, the LVs from 16 different dogs were segmented and combined to produce an average geometric model, which was meshed according to previous finite element mesh approaches^48^. This finite element model was then defined for implementation in FEBio, an open-source software tool for nonlinear finite element analysis specifically focused on solving large deformation problems in biomechanics and biophysics^49–51^. Since the MRI segments used to build this FE model were obtained at end systole, the fitted mesh was scaled to better approximate the measured EDV and unloaded LV geometry, which was taken to be stress-free in all simulations. This unloaded model had a cavity volume of 30 mL, and was used in Estrada’s optimization to obtain passive material parameters^46^. This Baseline model, representing a normal heart, was subsequently adjusted by Estrada and colleagues to additionally model Ischemic and Patched conditions.

It was from this Patched configuration that the FE model was further adapted in this study to evaluate the performance of an auxetic ventricular support device. The Estrada Baseline model comprised three distinct materials: apex, valve, and myocardium, with the apex and valve being represented as rigid bodies and the myocardium being represented as a transversely isotropic Mooney-Rivlin solid, with active contraction modeled using a custom formulation combining the length-dependent behavior currently used in FEBio and a modified force-velocity relationship based on the fading-memory formulation proposed by Hunter and colleagues^46,52^. For the Ischemia model, a subset of myocardium elements (2840 out of 7600 total) were redefined as ischemic tissue, also represented as a transverse isotropic Mooney-Rivlin solid, but with distinct material properties (and no active contraction) based on mechanical measurements obtained during previous *in vivo* studies^46^. For the Patch model, an epicardial subset of the elements (567 out of the 2840) that had comprised the ischemic region in the Ischemia model were redefined as patch (either isotropic, longitudinally reinforced, or circumferentially reinforced), which was also defined as a passive, transversely isotropic Mooney-Rivlin.

The validated Baseline, Ischemia and Isotropic Patch models published by Estrada and colleagues^46^ were used as they were obtained, with no modifications to their input files or boundary conditions. For modeling an implanted auxVSD, their Isotropic Patch model (herein, called the Meiotic Patch, and designated by a positive Poisson’s ratio, ν > 0) was modified to replace the original patch with a material having auxetic mechanical properties (Figure 1). Thanks to the material data structures used by FEBio, this swap was relatively straightforward and required only changing the material definitions for the patch subsection of the overall model material definition file. Indeed, the primary challenge in making this patch material swap was identifying a material definition that would allow the model to converge when the patch material had a negative Poisson’s ratio. Ultimately, a simple neo-Hookean definition was found to be compatible with the rest of the Patched FE models, allowing the explicit definition of patch material density, elastic modulus, and Poisson’s ratio.

**Figure 1.**
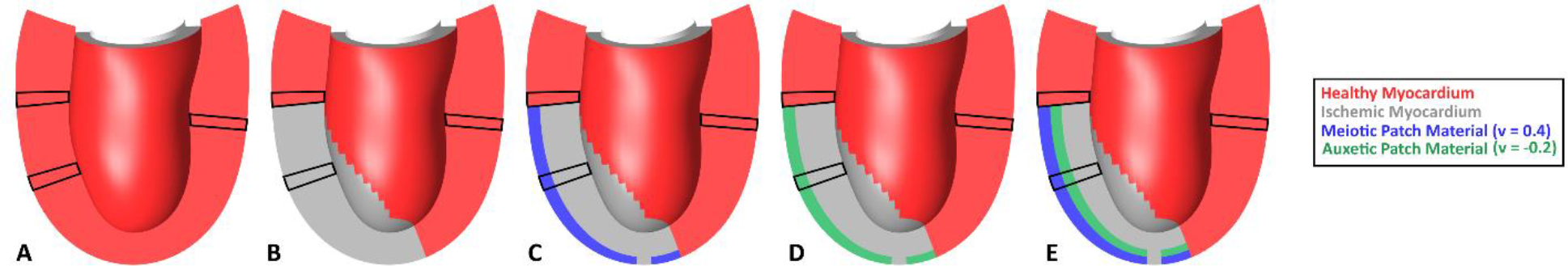
Schematic of each finite element model in the initial, unloaded state with the locations of the three sampling ROIs (patch+infarct, border zone, remote myocardium). (A) Baseline (B) Acute Ischemia (C) Meiotic Patch (D) Homogeneous Auxetic Patch (E) Heterogeneous Auxetic Patch

The Baseline, Ischemia, and Meiotic Patch models obtained from Dr. Estrada were all re-run locally and without any modification in FEBio v2.9.1 to confirm model convergence and results, which were identical to the previously published study^46^. For the homogeneous and heterogeneous auxVSD models, 10 patch variations were tested, varying the bulk elastic modulus of the patch from 10kPa to 100kPa by increments of 10kPa. For each variation, a density of 1.0g/mL was prescribed as a rough average for elastomer specific gravity^53^, and a Poisson’s ratio of -0.2 was prescribed, based on the average Poisson’s ratio measured during bench testing of some physical auxetic samples fabricated in our laboratory^54^.

For the purposes of this study three regions of interest were chosen for sampling simulation data from the LV computational model, representing key areas of physiological relevance: a remote myocardium region, a patch/infarct region, and a border zone region. Each region comprised a segment of five adjacent elements, traversing from the epicardium to the endocardium (Figure 1). The remote myocardium was taken as a segment of the healthy myocardium material directly opposite the elements defined as infarct/patch. In the case of the border zone region, the five-element segment included elements that were either part of the myocardium or part of the infarct/patch. Lastly, the infarct/patch ROI was in the middle of the region of elements in the LV model representing either a transmural segment through the Ischemic region, or a transmural segment including four elements of infarct and one (epicardial) element of patch in the Homogeneous Auxetic case, or three elements of infarct, one element of auxetic material and one element of meiotic epicardial patch material in the Heterogeneous Auxetic case. At each of these sampling ROIs, fiber stress and radial strain (relative to end diastole) over the cardiac cycle were compared between the Baseline, Ischemic, Meiotic Patch, and Auxetic Patch models. Additionally, P-V loops were generated for each of these models based on the change in cavity volume resulting from the prescribed pressure within the LV cavity, which was part of the original FE model and the same for all models tested^46^.

## Results

### Fiber Stress

The Baseline, Ischemia, and Meiotic Patch models all recapitulated the results found by Estrada et al. during initial testing of the model^46^. Unsurprisingly, fiber stress in the border zone ROI nearly doubled in the Ischemia model relative to Baseline, highlighting the mechanical conditions at the interface between still-contractile tissue and non-beating, mechanically compromised tissue. When the Meiotic Patch was added, though, the fiber stress in the border zone decreased to about 125% of baseline during systole, highlighting the mechanical offloading performed by the patch. Notably, the heterogenous patch reduced fiber stress in the border zone to about 65% of baseline levels. In the case of the homogeneous and heterogeneous Auxetic Patch models, elastic modulus was the primary determinant of patch performance, with low elastic modulus auxetic patches (10kPa to 30kPa) reducing the systolic fiber stress by a marginal amount, while high elastic modulus auxetic patches (80kPa to 100kPa) nearly matched the performance of the original Meiotic Patch Model (Figure 2). In the remote myocardium, systolic fiber stress remained roughly the same for all models tested (Figure 3); however, the patch conditions showed a slightly elevated diastolic fiber stress in the remote myocardium due to the greater ED volume in the ischemic and patched models compared to baseline, which was minimally affected by the patch design.

**Figure 2.**
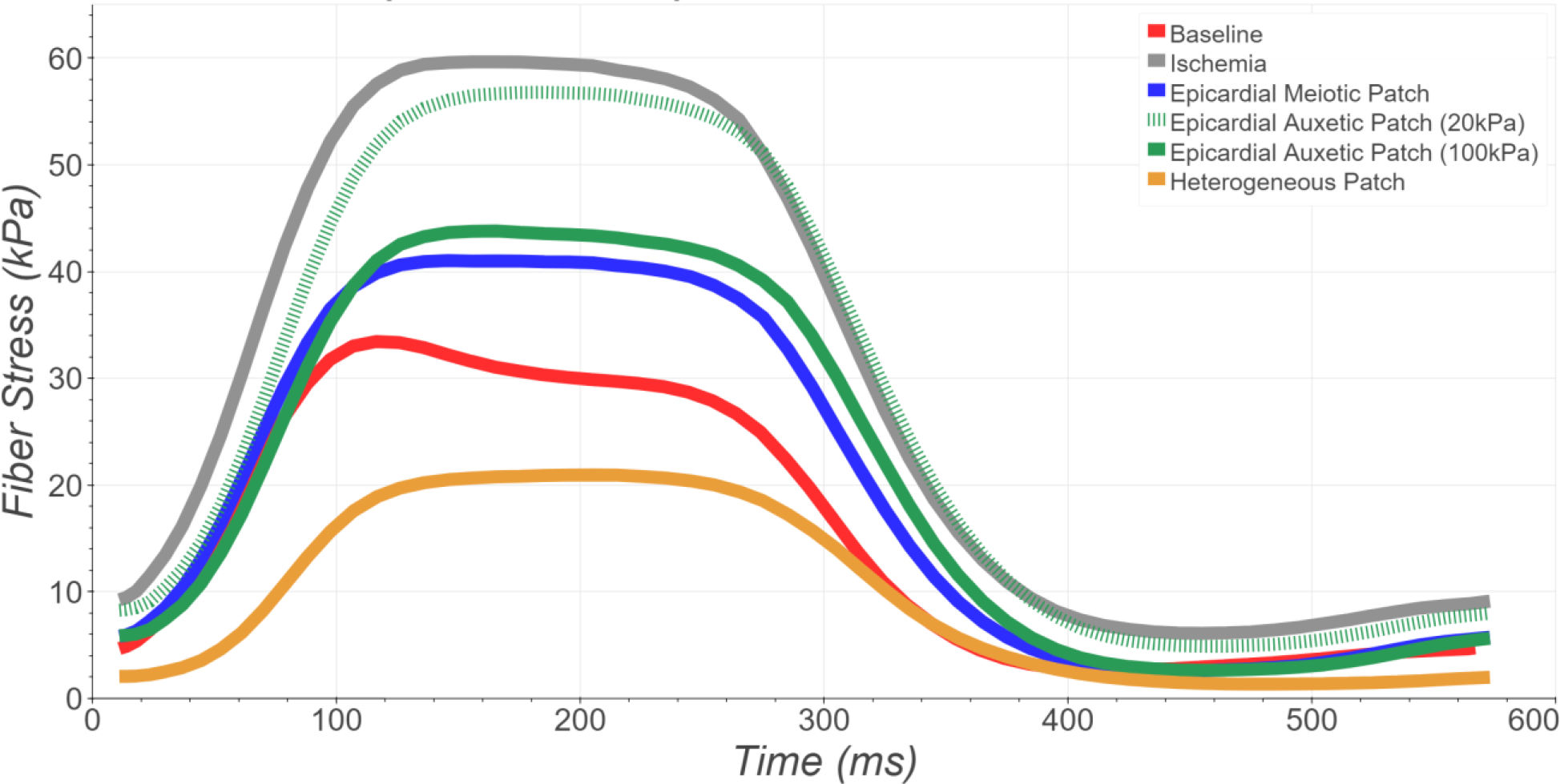
Border zone fiber stress from the mid-wall element, for Baseline, Ischemia, and all patched conditions. The low modulus homogeneous Auxetic Patch had little impact on fiber stress relative to the Ischemic condition; however, the higher modulus homogeneous Auxetic Patche reduced border zone fiber stress comparably to the Meiotic Patch condition, and the heterogeneous patch reduced it below baseline levels.

**Figure 3.**
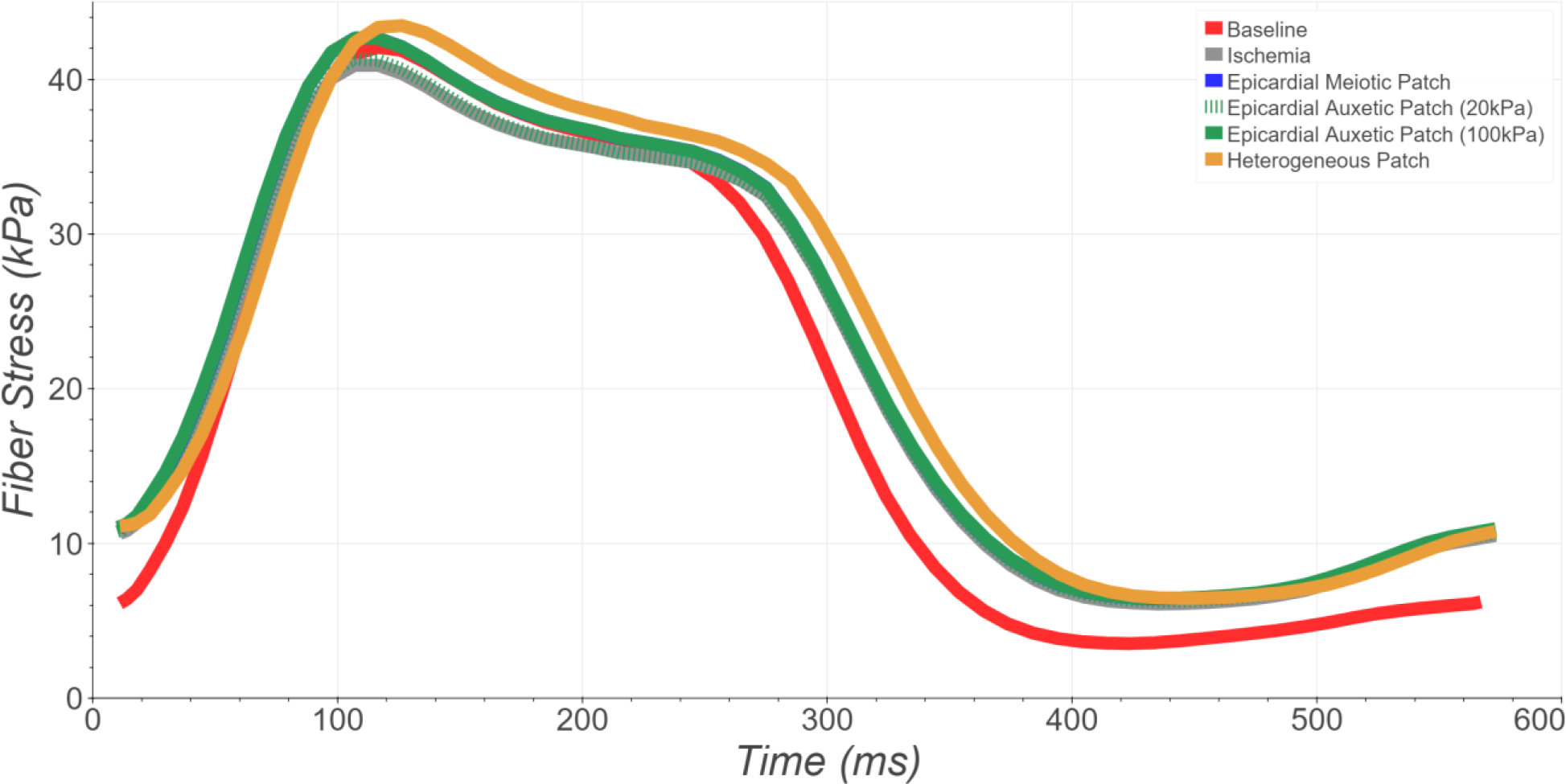
Fiber stress in remote myocardium, from the mid-wall element, for the Baseline, Ischemic, and all four patched conditions. All patched conditions yielded comparable results over the duration of the cardiac cycle, with diastolic stress slightly elevated compared to Baseline.

### Radial Strain

For radial strain, the ROI of primary interest was the patch/infarct region, running directly through the middle of those two materials in the FE model. Again, the original three models yielded expected behavior with the Baseline model exhibiting a positive radial strain, indicative of myocardial thickening during systolic contraction. In the case of the Ischemia model, this ROI exhibited negative radial strain throughout the entire systolic phase, indicative of thinning due to the local loss in contractility as well as stretching by the neighboring healthy myocardium. For the Meiotic Patch model, negative radial strain was also observed through the entire systolic phase, indicating thinning of the patch under pressure loading, as expected. Importantly, for both low and high elastic modulus homogeneous Auxetic Patch models, radial strain within the patch was positive due to the auxetic thickening caused by in-plane stretching of the patch during systole, achieving about a quarter of the strain magnitude of the Baseline model. Most impressive was the radial strain exhibited by the Heterogeneous Auxetic Patch, which achieved nearly half as much thickening as that exhibited by the healthy Baseline model (Figure 4), possibly indicating a mechanical benefit from interaction with the meiotic layer. These results provide encouraging evidence supporting the capacity of an auxVSD to undergo auxetic thickening driven by in-plane stretching from the neighboring healthy myocardium, which could partially restore local ventricular wall mechanics in the setting of acute MI.

**Figure 4.**
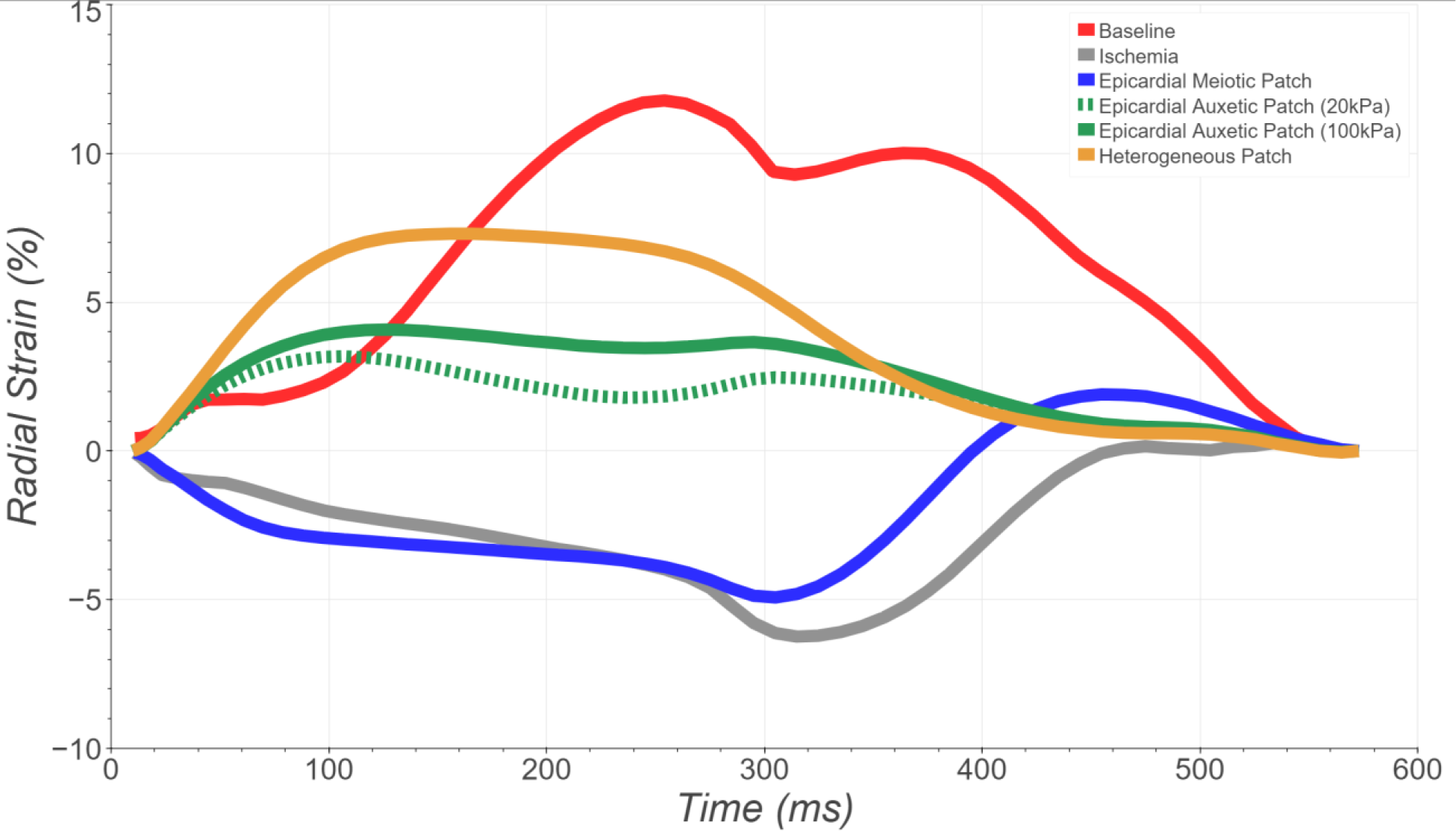
Radial strain in the patch/infarct region. demonstrates the systolic thickening of auxetic patches due to stretching by neighboring active myocardium. The Baseline model exhibited positive systolic radial strain (wall thickening), whereas the Ischemia and Meiotic Patch models exhibited negative radial strain (wall thinning). All three Auxetic Patch designs exhibited positive radial strain, indicative of auxetic expansion driven by the mechanical interaction with neighboring myocardium and underlying ischemic tissue causing the Auxetic Patch materials to stretch and thicken during systole. Strain measurements obtained from the epicardial element in each segment for all conditions except for the Heterogeneous Auxetic Patch, where the auxetic layer is one layer deeper than the other models.

### Pressure-Volume Loop Analysis

To understand how different patches impacted global cardiac mechanics and LV pump function, P-V loops were generated for the three initial models as well as the auxetic patch models (Figure 5). As expected, the Meiotic Patch was able to partially correct the post-ischemic increases in LV volume, lowering end-systolic volume (ESV) from approximately 63mL to 60mL, and end-diastolic volume (EDV) from 77mL to 74mL. The Meiotic Patch model also slightly increased ejection fraction (EF) to 18.9% versus 18.2% in the ischemia model. Comparing different Auxetic Patches, elastic modulus once again emerged as a strong driver of outcomes, with a low elastic modulus (20kPa) patch minimally altering LV mechanics relative to the Ischemia model. In the case of a high elastic modulus (100kPa) homogeneous Auxetic Patch, the P-V loop was very similar to that for the Meiotic Patch. However, the Heterogeneous Auxetic Patch model showed a much more pronounced leftward shift of the P-V loop, with ESV reaching 55mL, compared to 52mL from the Baseline model, and corresponding EDV values of 69mL and 67.5mL, respectively. Furthermore, EF was increased to 20.3% for the Heterogeneous Auxetic Patch model, approaching the value of 22.9% in the Baseline model. These findings support the potential for auxetic patch materials to achieve superior functional outcomes compared to VSDs created with traditional materials for treating post-MI ventricular dysfunction.

**Figure 5.**
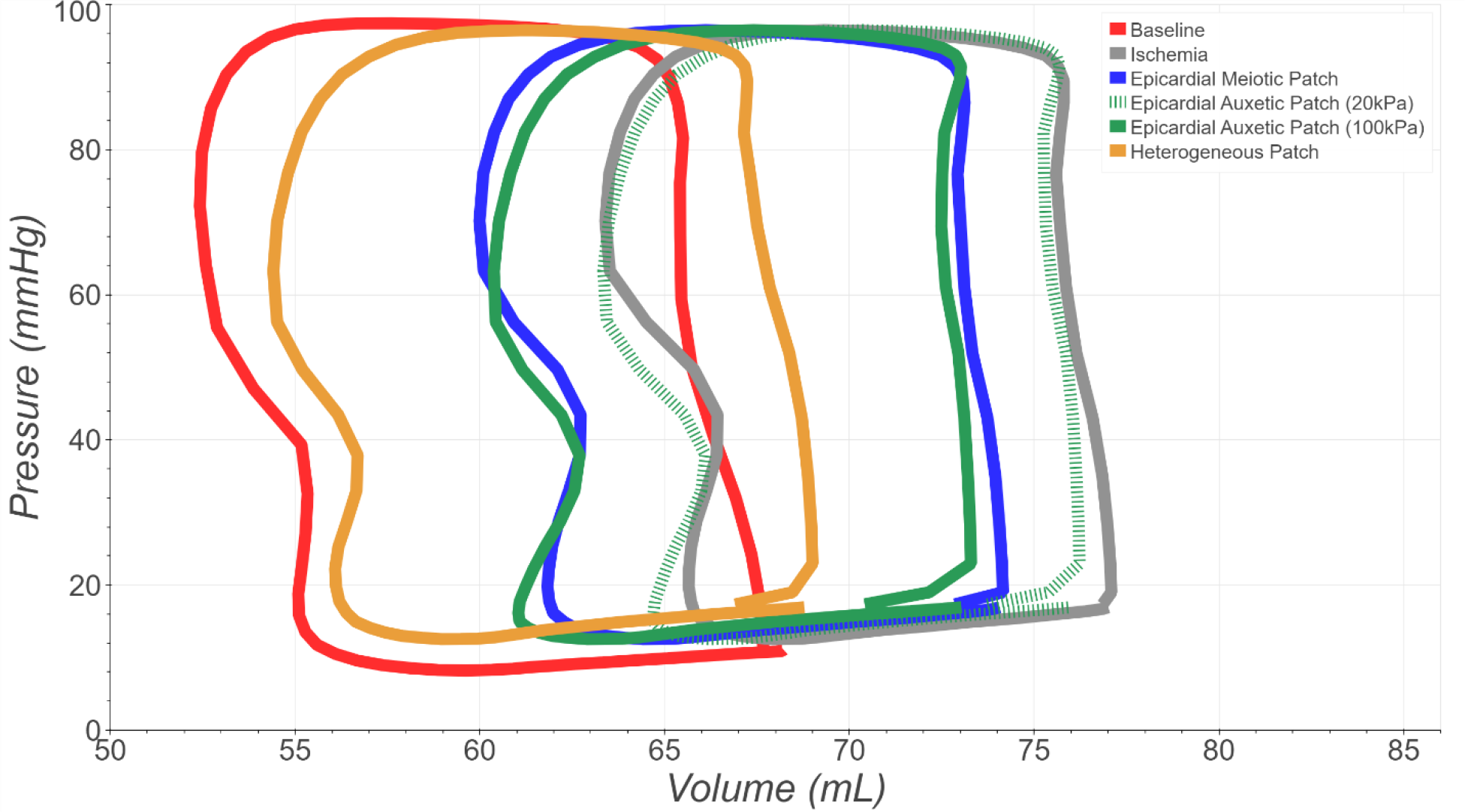
Left ventricular pressure-volume loops for all model conditions. Consistent with the fiber stress measurements, performance of the homogeneous Auxetic Patches were closely correlated to their respective elastic moduli, and not more effective than the standard Meiotic Patch. Dramatically improved performance of the Heterogeneous Auxetic Patch, however, indicates that introducing patch materials with directionally-biased mechanical behaviors can result in a more biomimetic performance.

## Discussion

The goal of this study was to explore the potential of utilizing an auxetic patch graft material for post-MI myocardial reinforcement that could dynamically respond to local cardiac mechanics and yield better global cardiac performance than a traditional, meiotic patch graft material. Leveraging the previously-validated biomechanical model of canine left ventricle from Estrada and colleagues^46^, we were able to model this scenario under several different conditions and auxetic metamaterial patch designs. Overall, these simulations yielded encouraging preliminary results and provide a strong starting point for future modeling studies, patch design refinement, and *in vivo* experiments. As evidenced by local radial strain results within the patch/infarct material, auxetic activation and thickening of both homogeneous and heterogeneous patch designs did occur, driven by the stretching forces imparted by adjacent, actively contracting myocardial elements. Furthermore, the auxetic thickening at the patch/infarct region did not appear to dramatically alter the stresses in remote myocardium, despite slightly elevated diastolic stresses, indicating minimal “off-target” mechanical consequences in other regions of the LV.

At the whole-ventricle level, the impact of an auxetic patch graft was more mixed, with homogeneous auxetic patches either underperforming (low elastic modulus) or essentially matching (high elastic modulus) the performance of the Meiotic Patch, with respect to the P-V relationship. It may be that this lack of global impact despite local auxetic thickening was due in part to the unconstrained bi-directional radial expansion of the auxetic patch, with some potential benefits of the auxetic effect being lost. Further supporting this hypothesis were the results of the Heterogeneous Auxetic Patch model, which most closely resembled the mechanics of the Baseline model with its P-V loop shifted farthest to the left, and the biggest improvement in ejection fraction, among all tested patch conditions. This pronounced difference in global cardiac mechanics between the stiff homogeneous and heterogeneous auxetic patch graft models suggests that, with the proper design, an auxetic patch graft can be superior in performance to traditional patch graft materials, with plenty of opportunity for subsequent improvements to patch design.

Indeed, the isotropic linear elastic patch material used in this study is perhaps the simplest but least potent form of an auxetic patch. It is possible that greater efficacy can be achieved with hyper-auxetic metamaterials that have large negative values of Poisson’s ratio^55–57^, and anisotropic behavior that could maximize radial thickening beyond the current scenario. This is theoretically possible using an orthotropic elastic material model with negative Poisson’s ratio exceeding -1, but such materials are not yet implemented in FEBio. In practice, advanced manufacturing techniques such as 3D printing and CNC knitting could be used to fabricate metamaterials with such specialized properties. It may also be possible to fabricate auxetic metamaterials with built-in structural heterogeneity, so that it would not be necessary to combine auxetic and meiotic layers in order to bias the direction of auxetic expansion. Such advancements represent exciting areas for future investigation.

## Limitations

One limitation of this study is that the same LV pressure loading timecourse was used for all models, when in fact the LV pressure would be expected to differ in healthy vs. post-infarct conditions. However, this allowed us to more directly compare the potential effects of each patch material on LV mechanical function, and in the absence of experimental data to validate each case, this was considered a reasonable starting point. Another limitation is that finite element models of LV pump function are typically not as efficient as the native heart^58,59^, with the baseline model achieving an ejection fraction of only 20.3%. Therefore, validation of the computational model predictions using experimental animal models will be necessary to confirm the trends revealed in this study. Nevertheless, the relative changes in pump function showed clear advantages of the reinforced auxetic patch for restoring chamber function post-infarction, with biased auxetic thickening improving ejection compared to the other patch conditions. Considering the conservative auxetic properties assigned to these patch materials (ν = -0.2), it seems likely that with additional optimization, metamaterials with a more strongly hyper-auxetic behavior could offer further improvements with potential for clinically significant outcomes for patients with post-infarction heart failure. In this context, it is also worth noting that the model of acute ischemia may not represent the most relevant clinical scenario for patch VSD reinforcement; a more urgent clinical need would be patients with large chronic infarction with transmural scar tissue that is expanding and forming a dyskinetic aneurysm. However, such a finite element model has not yet been developed and experimentally validated as a starting point to test the mechanics of auxetic patch plasty.

## Conclusions

Although a homogeneous auxetic patch did not appear to dramatically alter global cardiac mechanics relative to a meiotic patch due to the lack of constraint/bias on the direction of auxetic expansion, favorable local mechanics observed during initial simulations was able to guide the development of a heterogeneous patch model with mechanical performance much closer to Baseline. Furthermore, while the homogeneous auxetic patches showed no meaningful improvement over meiotic patches at the global level, their local mechanical behavior at the infarct/patch ROI, as well as the performance of the heterogeneous patch, all clearly demonstrate that an auxVSD patch coupled to remote myocardium can indeed harness sufficient stretching force to expand against an LV pressure load. Lastly, the general success of the heterogeneous patch at restoring key features of cardiac mechanics toward the Baseline condition demonstrates how this computational model served as a virtual screening platform that was able to inform the development of an improved patch design. Further experimental studies will be required to validate whether such auxetic patch designs can be applied to achieve similar functional improvements *in vivo*.

